# Honeybees collect pollen from the buzz-pollinated flowers of invasive *Solanum elaeagnifolium* in Northern Greece

**DOI:** 10.1101/2025.05.13.648936

**Authors:** A. Kantsa, G. Nakas, R.R. Kariyat, H. Lambert, T. Petanidou, C.M. De Moraes, M.C. Mescher

**Affiliations:** Department of Environmental Systems Science, ETH Zurich, Switzerland; Department of Geography, University of the Aegean, Greece; Department of Entomology C Plant Pathology, University of Arkansas, Fayetteville AR, USA

**Keywords:** Agroecology, *Apis mellifera*, Buzz-pollination, Plant invasions, Pollen resources, Solanaceae, Weed ecology

## Abstract

Around 6% of flowering plants, including most solanaceous species, have flowers specialized for buzz-pollination (sonication). The poricidal anthers of these flowers release pollen in response to frequency-specific vibrations, which buzz-pollinating bees, including bumble bees and many solitary bees, generate via rapid oscillation of flight muscles. Honeybees are incapable of sonication and typically avoid visiting buzz-pollinated plants, although they have previously been observed to scavenge residual pollen from petals of buzz-pollinated flowers in natural settings. Here, we report a novel interaction in which honeybees consistently visit and extract pollen from flowers of *Solanum elaeagnifolium* (silverleaf nightshade) within a large invasive plant population in Northern Greece. We postulate that honeybee foraging effort on *S. elaeagnifolium* reflects the shortage of alternative floral resources at the peak of the Mediterranean summer, consistent with observations for pollen-starved bees in captivity. This newly established behavioral shift of honeybees may convey important implications for plant invasions, particularly under climate change, and requires further investigation for untangling its ultimate drivers.

## Introduction

> “…the polylectic honeybee does not seem able to learn the process.”
>
> Linsley (1978)

Buzz-pollinated flowers occur in around 20,000 species across various angiosperm clades (∼6% of species), including key crops such as tomato, eggplant, blueberry, cranberry, and kiwi (Buchmann 1983, De Luca and Vallejo-Marín 2013, Pritchard and Vallejo-Marín 2020, Cooley and Vallejo-Marín 2021). Owing to their unique floral morphology, the poricidal anthers of these flowers require vibration at specific frequencies (also known as sonication) to release pollen. Buzz pollination is hypothesized to serve as an adaptation for preventing pollen access by relatively inefficient, non-specialist pollinators (Pritchard and Vallejo-Marín 2020), although the morphology of buzz-pollinated flowers may also deter herbivory by making pollen less accessible (Hargreaves et al. 2009). A capacity for sonication via rapid oscillation of the thoracic indirect flight muscles has evolved independently ca. 45 times within bees (Cardinal et al. 2018), enabling approximately 58% of bee species (at least 74 genera) to efficiently extract pollen from buzz-pollinated flowers (Buchmann 1983, De Luca and Vallejo-Marín 2013, Pritchard and Vallejo-Marín 2020, Jankauski et al. 2022). However, honeybees, the most frequent floral visitors in both natural and agricultural systems worldwide (Hung et al. 2018), are incapable of sonication, and their limited interaction with buzz-pollinated plant species has been widely discussed by pollination biologists (Linsley 1978, Buchmann 1983, De Luca and Vallejo-Marín 2013, Vallejo-Marin et al. 2024).

The proximate mechanisms preventing honeybees from effectively collecting pollen from buzz-pollinated flowers appear to be based in physiological constraints—specifically relating to the maximum sustained frequency of thoracic vibrations (King and Buchmann 2003)—rather than a lack of behavioral flexibility or learning capacity, as had previously been suggested (Linsley 1978). In any case, honeybees typically avoid buzz-pollinated flowers in natural settings, and particularly those of species that do not offer nectar, as in the genus *Solanum* (Michener 1962, Linsley 1978, Buchmann 1983, Buchmann and Cane 1989, Tscheulin et al. 2009, Tscheulin and Petanidou 2013, Petanidou et al. 2018, Tayal and Kariyat 2021). Οn rare occasions in which honeybees have been observed to visit buzz-pollinated flowers in the wild, they have primarily been reported to glean leftover pollen from petals following visits by buzz-pollinators (Wille 1963, Buchmann 1983, Buchmann and Cane 1989, Gross and Mackay 1998, Solís-Montero et al. 2015, Mesquita-Neto et al. 2018, Petanidou et al. 2018, Rocha et al. 2023), and sometimes to steal already deposited pollen from the stigma, presumably reducing plant fitness (e.g., Gross and Mackay 1998).

Despite their limited ability to utilize buzz-pollinated flowers, honeybees can be induced to visit such flowers under conditions of pollen deprivation and are sometimes utilized for pollination of buzz-pollinated agricultural crops. In the case of buzz-pollinated flowers that do not offer nectar (e.g., tomato, eggplant, potato, tamarillo), managed honeybees will visit only when confined into greenhouses or flight cages with no access to other pollen sources (Neiswander 1956, Cane et al. 1993, Cane and Schiffhauer 2003, Miyamoto et al. 2006). However, honeybees will spontaneously visit only the nectar-offering buzz-pollinated crops (e.g., blueberry, cranberry), in which case they can be used in open environments (Cane et al. 1993, Dogterom and Winston 1999). The reported pollinating efficiency of honeybees on buzz-pollinated crops ranges from zero (avoided the flowers, although pollen-deprived; e.g, Sanford and Hanneman 1981, Banda and Paxton 1990, Amala and Shivalingaswamy 2017, Zhang et al. 2020, Rijo et al. 2022), to inconsistent (Higo et al. 2004, Dos Santos et al. 2009, Courcelles et al. 2013, Cooley and Vallejo-Marín 2021), or sometimes satisfactory provided that hives are used in high densities and honeybee workers do not steal nectar (Neiswander 1956, Cribb et al. 1993, MacKenzie 1994, Delaplane and Mayer 2000, Cane and Schiffhauer 2003, Sabara and Winston 2003, Miyamoto et al. 2006, Courcelles et al. 2013, Cooley and Vallejo-Marín 2021).

The observation that managed honeybees will visit and attempt to collect pollen from buzz-pollinated flowers under pollen deprivation, raises the possibility that similar behaviors may occur in nature, particularly in environments where honeybees have limited access to other pollen sources or, conversely, where other potential pollinators are absent or rare. Both conditions might arise in the context of plant invasion (Albrecht et al. 2016, Davis et al. 2018).

Yet, as noted above, reports of honeybee visitation of buzz-pollinated flowers in nature are rare and generally limited to observations of bees collecting residual pollen from petals, following visitation by other, buzz-pollinating species. In the current study, we report the large-scale extraction and utilization by honeybees of pollen from a buzz-pollinated plant (*Solanum elaeagnifolium*), which forms large monocultures in its invaded (Mediterranean) range and produces flowers in a seasonal period when other flowers are scarce. We also provide evidence that this honeybee behavior has become widespread within the last decade and discuss potential ecological and evolutionary implications within the context of plant invasion.

## Materials and Methods

### Study species

*Solanum elaeagnifolium* (silverleaf nightshade) is currently among the top-noxious invasive plants worldwide (Uludag et al. 2016, Roberts and Florentine 2022). A perennial herbaceous species originating in N Mexico and SW USA, *S. elaeagnifolium* has been rapidly expanding to almost all continents over the past century, posing serious threats to native biodiversity and causing significant losses in agricultural and livestock production in many countries (Uludag et al. 2016, Roberts and Florentine 2022). Its uncontrollable expansion has severely affected the Mediterranean Basin, making it one of the most impacted regions (Uludag et al. 2016, Krigas et al. 2021). In croplands and urban areas of N Greece, this species forms numerous extensive monospecific stands (of >10,000 individuals) (Petanidou et al. 2018) (Figure 1).

**Figure 1.**
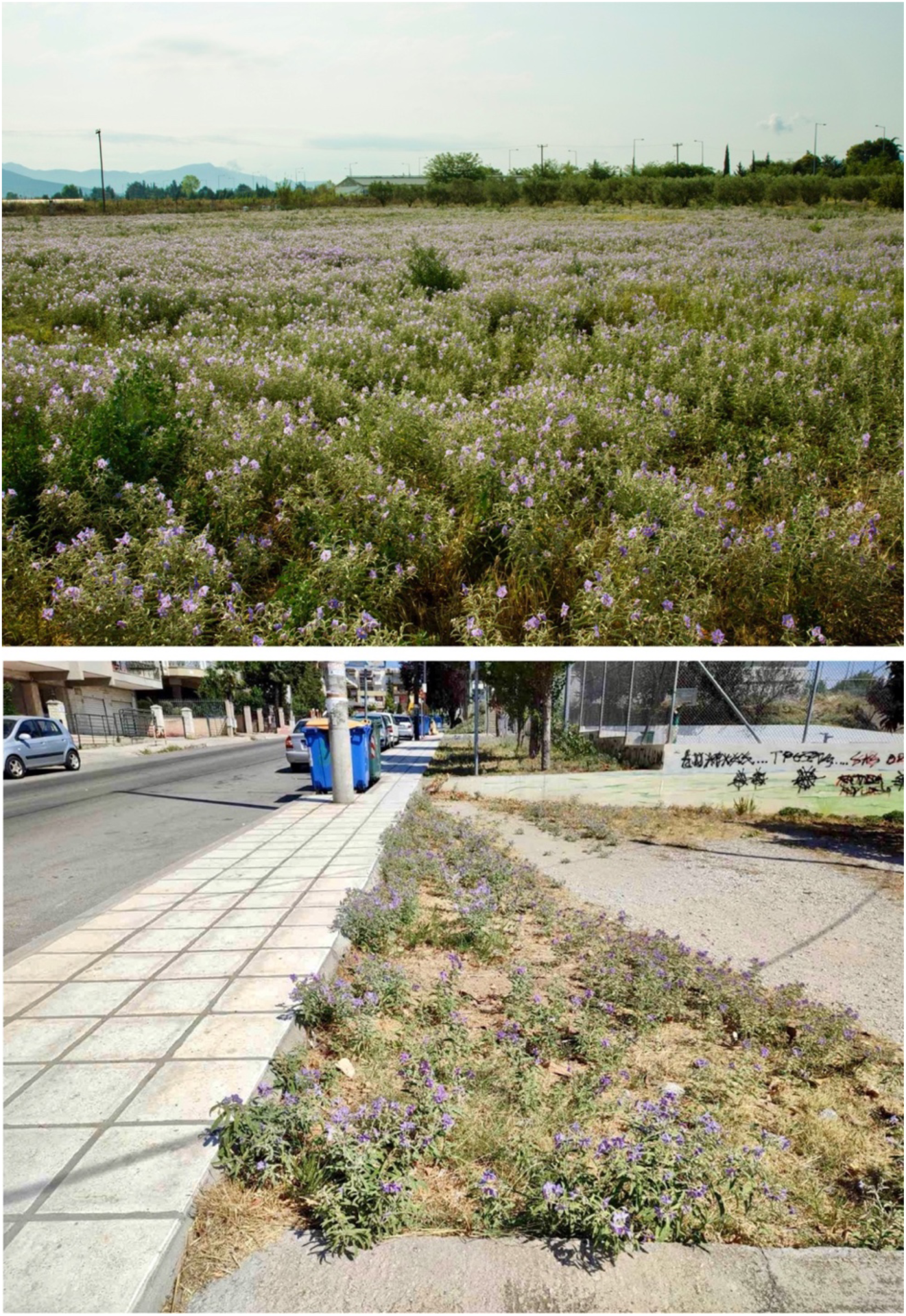
Invasive population stands in a cultivated field (up) and in an urban residential area (down) in the Metropolitan area of Thessaloniki, N Greece (photos: R.R. Kariyat, G. Nakas).

### Bee observations

Bee observations on *S. elaeagnifolium* were performed in July-August 2015, 2016, 2017, and 2023 in the Metropolitan Area of Thessaloniki, Greece (Table 1; Figure 1), at sites with either urban or agricultural land use, which had been monitored in the context of previous research of this species in the area in 2006-2007 (Petanidou et al. 2012, Petanidou et al. 2018). Observations were made during 30-min random walks across each study site in the morning hours (09:00-12:00). Honeybees that visited the flowers and tried to extract pollen using any of the behaviors described above were photographed and video-recorded, and samples were brought to the lab in order to identify the pollen they carried.

**Table 1.** Study sites, surveys, and honeybee observations on the flowers of *Solanum elaeagnifolium* in Thessaloniki. The years of the first and the last surveys performed at each site, as well as the year in which the interaction between the plant and honeybees was first observed are shown.

| Site ID | Site name | Coordinates | Land use | First survey | Last survey | First observation |
| --- | --- | --- | --- | --- | --- | --- |
| S01 | Sindos | 40° 41' 6.0" N,<br>22° 43' 40.8" E | Agricultural | 2007 | 2016 | - |
| S02 | Technological Institute | 40° 40' 12.0" N,<br>22° 47' 20.4" E | Agricultural | 2007 | 2023 | - |
| S03 | Anchialos | 40° 43' 32.2" N,<br>22° 41' 39.1" E | Waste land | 2007 | 2016 | - |
| S04 | Redaistos | 40° 30' 58.88" N,<br>23° 4' 21.37" E | Agricultural | 2007 | 2017 | 2015 |
| S05 | Panorama | 40° 35' 26.72" N,<br>23° 2' 38.96" E | Semi-natural | 2007 | 2015 | - |
| S06 | Asvestochori | 40° 40' 15.6" N,<br>23° 3' 0.0" E | Semi-natural | 2007 | 2015 | - |
| S07 | Thermi | 40° 32' 6.29" N,<br>22° 59' 45.12" E | Agricultural | 2016 | 2023 | 2017 |
| S08 | Airport | 40° 31' 10.9" N,<br>22° 58' 15.2" E | Semi-natural | 2023 | 2023 | 2023 |
| S09 | City center | 40° 37' 20.22" N,<br>22° 57' 17.89" E | Urban parks | 2023 | 2023 | 2023 |
| S10 | Stavroupoli | 40° 39' 32.8" N,<br>22° 55' 8.8" E | Urban parks | 2023 | 2023 | - |

### Scanning electron microscopy (SEM)

Honeybees foraging on *S. elaeagnifolium* were captured and brought to the lab to confirm the presence of pollen from this plant in their corbiculae and on their bodies. For this, we compared SEM images of the pollen of this species and of the pollen found on the honeybee specimens [see also Du et al. (2018)]. SEM was performed at the Scientific Center for Optical and Electron Microscopy of ETH Zurich (ScopeM), using a multi-purpose analytical high resolution field emission gun scanning electron microscope (HR-FEG-SEM Quanta200F). Images were acquired at a 10 kV accelerating voltage under low vacuum conditions (100 Pa), with the secondary electron (SE) signal recorded using the LFD detector. The field of view (FOV) was 149 µm or 298 µm, depending on the imaging setup. Since charging was suppressed by using the low vacuum mode, no metal coating was applied on the bees before introducing them to the SEM vacuum chamber (Symondson and Williams 1997).

## Results and Discussion

Since 2015, beginning in the E–SE part of the Thessaloniki metropolitan area and gradually expanding towards the city center (Figure 2), we have frequently observed honeybees visiting *S. elaeagnifolium* flowers and actively extracting pollen from the poricidal anthers using three distinct behaviors: (a) shaking/drumming the anthers using their forelegs; (b) biting off pieces of anthers with their mandibles; (c) holding the anthers with their forelegs and inserting their proboscis through the pores of the anthers (Videos S1-S3). As a result, pollen of silverleaf nightshade adheres on the bodies of honeybees and is used to fill up the corbiculae (Videos S1-S3) (Figure 3). These modes of behavior have previously been reported for honeybees trying to exploit buzz-pollinated flowers in cultivation, and Cane et al. (1993) hypothesized that the foreleg drumming behavior is a permutation of pollen-grooming movements that bees perform to transfer pollen grains from their upper body parts to the corbiculae.

**Figure 2.**
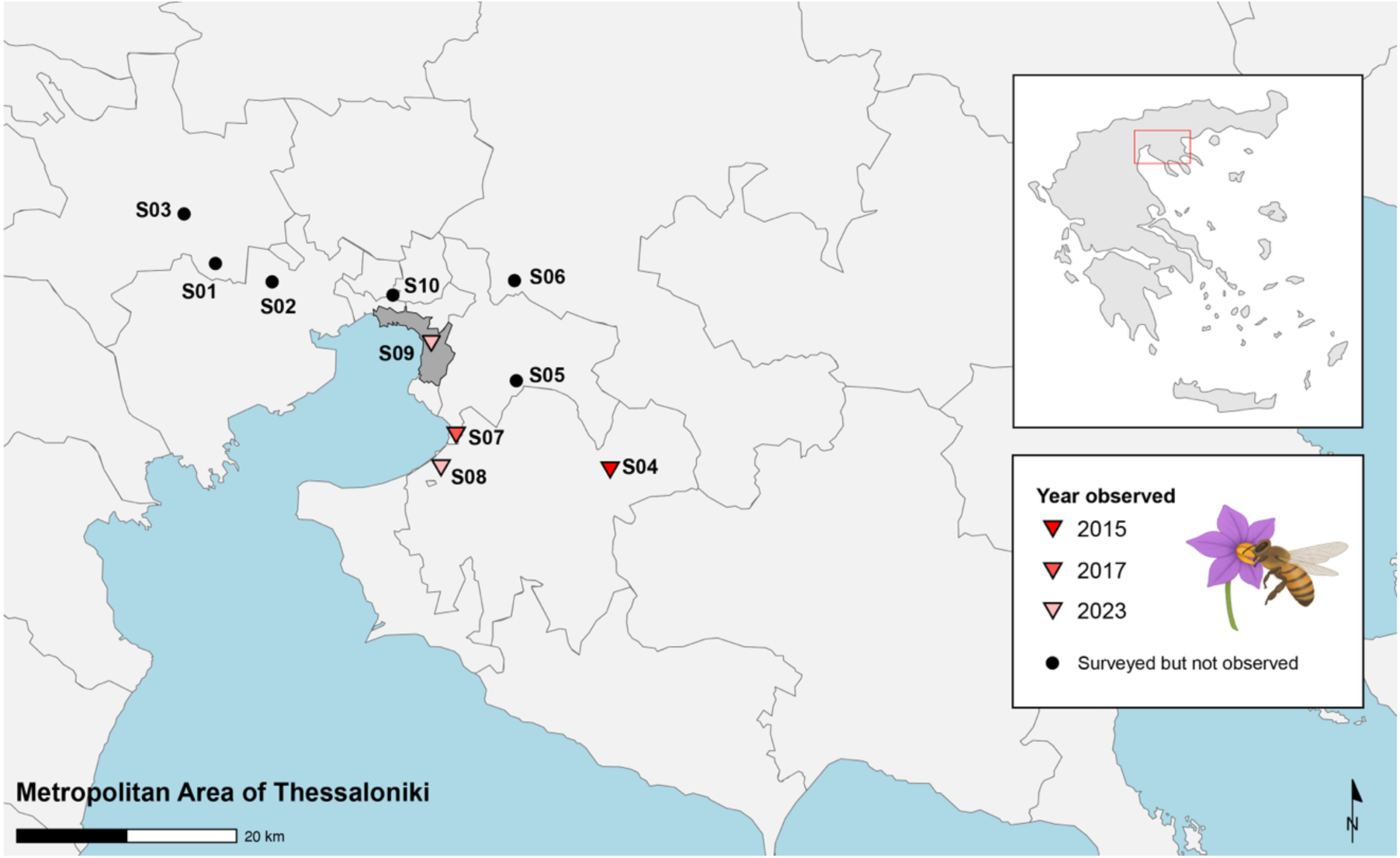
Study sites of *Solanum elaegnifolium* in the Metropolitan Area of Thessaloniki. In the sites where the honeybees have been observed to interact with the flowers, the year of the first observation is shown. The city center is marked with dark grey. See Table 1 for details on the sites.

**Figure 3.**
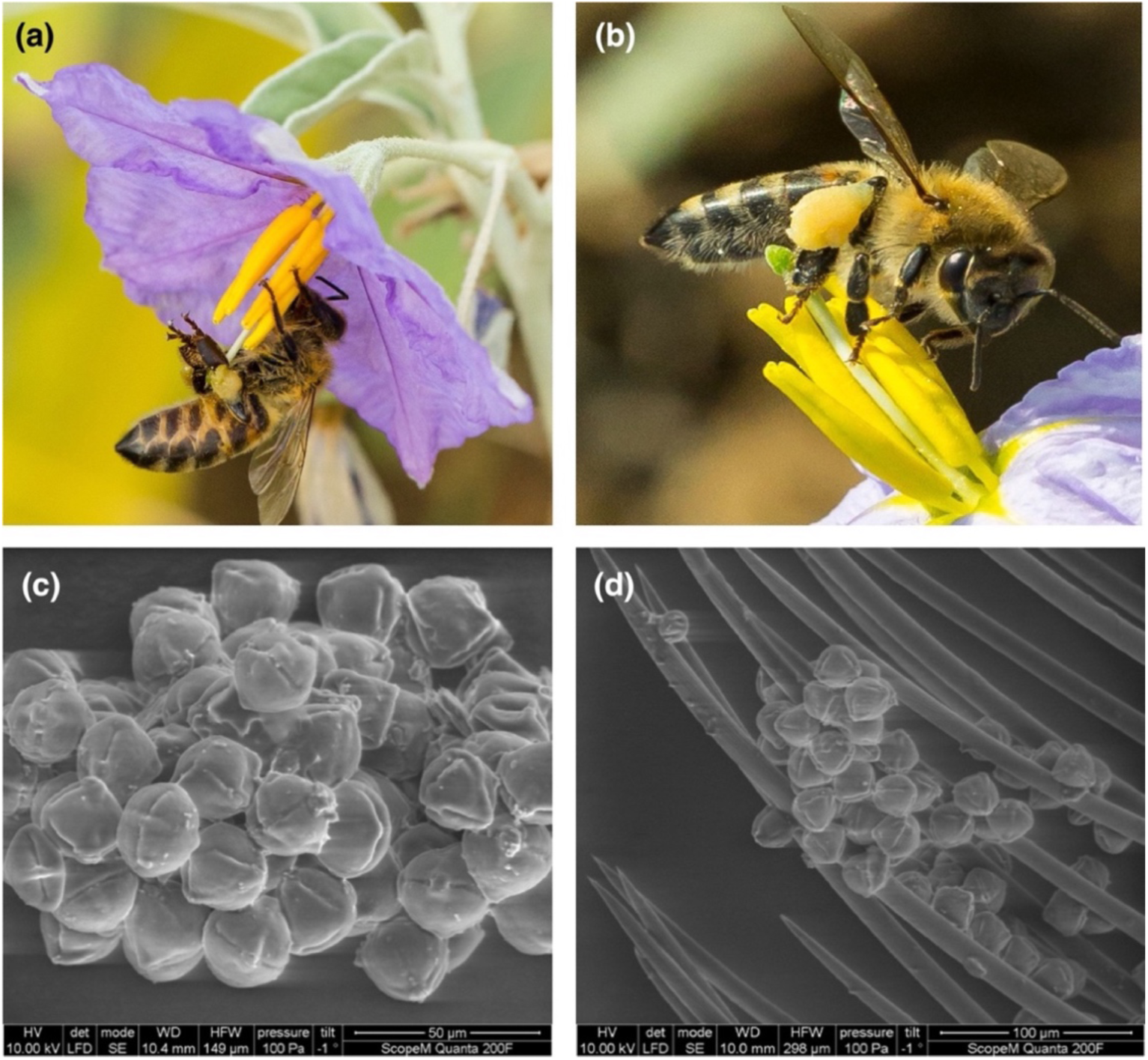
(a-b) Honeybees foraging for pollen on *Solanum elaeagnifolium*; their corbiculae (pollen baskets) are replete with the pollen of the plant (photos: R.R. Kariyat). (c-d) SEM images showing the pollen grains, and how these accumulate among the bristles on the body of the bee.

As documented in agricultural settings, honeybees spend more time on visitation of buzz-pollinated flowers than they usually do on other types of flowers, presumably due to the low efficiency of pollen extraction (Kendall et al. 2022). However, honeybee interactions with *S. elaeagnifolium* entail a high level of settings, physical effort and an exceptionally long-duration of floral visits. Indeed, in Thessaloniki, we recorded visits on single flowers lasting as long as 224 s (Video S1), whereas known maximum visitation times of honeybees on buzz-pollinated species are 100 s, observed on blueberry flowers (Kendall et al. 2022). Instead, for non-buzz-pollinated species, handling times are considerably shorter, typically ranging from 2.6 s per flower (oregano) and 4 s (lavender) (Hennessy et al. 2021) to 40 s (melon) (Ribeiro et al. 2017).

Furthermore, especially during the latter part of the flowering season (August), honeybees in Thessaloniki regularly visited *S. elaeagnifolium* flowers that were in poor condition, (e.g., heavily damaged by florivores, wilting, or deformed), which otherwise is known to reduce attractiveness to pollinators (e.g., Sõber et al. 2010, Vega-Polanco et al. 2020). Finally, it should be noted that honeybees are now by far the most frequent visitors of these flowers, outnumbering buzz-pollinating bees in the area (i.e., *Bombus, Pseudapis, Xylocopa,* and *Amegilla*) (RRK and GN personal observations). To our knowledge, this is the first report of honeybees consistently extracting pollen from a buzz-pollinated plant in a natural community.

The striking similarity of the behavior reported here with that previously reported for pollen-starved honeybees in confined agricultural settings (see above), suggests that it might be linked to severe limitation of other pollen sources. This is highly plausible given the magnitude of the invasion of silverleaf nightshade in the study area (Petanidou et al. 2018). It has been estimated that *S. elaeagnifolium* reached current population sizes in and around Thessaloniki city within 50 years after its first unintentional introduction (most probably from Texas) in the 1930s, forming one of the most heavily invaded areas in Europe (Krigas and Kokkini 2004, Uludag et al. 2016, Roberts and Florentine 2022). While we cannot be certain about when the widespread utilization of *S. elaeagnifolium* flowers by honeybees in Thessaloniki began, we can be confident that it started later than 2007, based on previous extensive surveys of pollinator activity in this area (Petanidou et al. 2012, Petanidou et al. 2018). While these surveys reported that other wild bees were scarce (Petanidou et al. 2018), honeybee interactions with *S. elaeagnifolium* were also rare, and restricted to gleaning leftover pollen from petals, presumably following visits by other pollinators (Petanidou et al. 2012, Petanidou et al. 2018). Similar behavior has been repeatedly reported for silverleaf nightshade both in Greece (Tscheulin et al. 2009, Tscheulin and Petanidou 2013) and in its native range, i.e., Texas and Arizona (Buchmann and Cane 1989, Tayal and Kariyat 2021). Besides, this fits the typical interaction of honeybees with *Solanum-*type flowers in the wild, where they have been reported to either avoid (*S. wedlandii*) (Michener 1962), or glean pollen leftovers (*S. rostratum*) (Solís-Montero et al. 2015) from flowers of *Solanum* species.

Since the successful establishment of silverleaf nightshade in Thessaloniki, local honeybee populations have presumably been exposed to a novel “operative environment” (Spomer 1973, Agosta and Klemens 2008), including a super-abundant pollen source, which, despite being challenging to exploit, is available during the most adverse time of year. Given the relative scarcity of other insect-pollinated species in flower in the lowlands of the Mediterranean at the peak of the dry summer (Kantsa et al. 2023b), silverleaf nightshade is by far the dominant pollen source in the invaded habitats. Therefore, we assume that the observed utilization of *S. elaeagnifolium* is a consequence of plant invasion in an area with limited alternative pollen sources, although it remains unclear why the behavior appears to have become widespread only in recent years.

The well-known aversion of honeybees towards buzz-pollinated flowers suggests that the utilization of nectarless, buzz-pollinated flowers is not adaptive in typical environments, likely because the ratio of pollen rewards obtained to effort expended is unattractive, relative to that provided by other flower types (Buchmann and Cane 1989). If utilization of *S. elaeagnifolium* flowers has been adaptive under the novel conditions in Thessaloniki, a question is whether this arises due to pre-existing behavioral flexibility or entails genetic adaptation. The apparently similar exploitation of buzz-pollinated flowers by pollen-starved honeybees in agricultural settings suggests that the former is plausible; the employment of different techniques by the honeybees (viz., drumming, “milking”, or biting; Videos S1-S3) is consistent with ecological fitting (Janzen 1985, Agosta and Klemens 2008), suggesting that the eventual mode of action depends on individual decision-making ability, agility, or both. These behaviors may thus represent a notable example of behavioral plasticity of the honeybees, facilitating exploration of novel areas of fitness space created by the massive invasive populations of *S. elaeagnifolium* in Northern Greece.

Regarding why the reported behavior was not observed prior to 2015, it may be useful to explore the potential preference of relevant colony-level heritable traits (e.g., pollen hoarding) in the local beekeeping market. For instance, Cane and Schiffhauer (2001, 2003) suggested that when honeybee queens are genetically predisposed for pollen hoarding (Rueppell 2014) pollen-foraging trips on buzz-pollinating flowers are significantly more frequent. Furthermore, the previously mentioned hypothesis by Cane et al. (1993) that the movement of foreleg drumming in honeybees is a permutation of pollen-grooming movements that bees perform to transfer pollen grains from their upper body parts to the corbiculae, suggests the possibility of an existing pre-adaptation. This could indicate a case of exaptation, where a trait originally evolved for one function (pollen grooming) has been co-opted for another (buzz-pollination). A similar example is the woodpecker finches (*Camarhynchus pallidus*, Thraupidae) at the Galápagos Islands, in which beak morphology originally suited for cracking seeds or eating insects has been co-opted for tool utilization, a behavior which varies according to habitat type (i.e., operative environment), is not acquired with social learning (but possibly with trial-and-error learning at a young age), and is believed to be based on genetic predisposition (Tebbich et al. 2001). In any case, whether honeybees have established a novel interaction circumventing adaptive processes (through phenotypic plasticity), or we are witnessing the real-time establishment of an exapted behavior, possibly conveying a more subtle and complex genetic background (e.g., pleiotropic traits), this phenomenon requires in-depth investigation.

Honeybees are globally abundant, frequently utilized in agriculture, and highly competitive with wild bees, often with adverse effects on native bee faunae (Goulson 2003, Lázaro et al. 2021). Given that they are generally among the first pollinators to interact with newly introduced plants in natural communities (Kantsa et al. 2023a), these behavioral shifts can be directly linked with invasion status and dynamics, with *S. elaeagnifolium* providing a characteristic example. In general, the rapid geographical expansion of this species is thought to be promoted largely by its ability to propagate vegetatively with root fragments (Roberts and Florentine 2022). The invasive populations of silverleaf nightshade are known to be self-incompatible, and (at least until 2007) pollen-limited (Petanidou et al. 2012, Petanidou et al. 2018). Based on these observations, it had been suggested that sexual reproduction was less important for the expansion of this plant in the invaded area. Yet, if this condition is overturned, and honeybee visitation can now convey the advantages of outcrossing, the genetic robustness of this vast invasive population is expected to be even further enhanced, with potentially even more disastrous effects on the native wildlife and landscape. For example, in the genus *Solanum,* outcrossing maintains high chemodiversity of volatile emissions, and it relates to better performance in chemically repelling herbivores and in recruiting predators (Kariyat et al. 2012a, Kariyat et al. 2012b, Kariyat et al. 2013).

## Outlook

Here we report a new foraging behavior in honeybees, which has emerged during the past decade, probably owing to the substantial expansion of *S. elaeagnifolium*, resulting in an overabundance of pollen during the most adverse period of the year in the Mediterranean in terms of floral resources. We have argued that our observation, likely involving a complex interplay of honeybee biology and floral ecology, has important implications both for the invasion status of silverleaf nightshade and for honeybee biology.

The enormous invasive populations of silverleaf nightshade in Northern Greece (and probably elsewhere) in fact represent large-scale natural experiments, which can reveal fascinating aspects of the evolution of plant–pollinator interactions. For instance, we know that buzz-pollination is not a dead-end, and novel floral forms (that do not even require bees) can evolve from bee-buzz-pollinated flowers (Dellinger et al. 2019). In the system examined here, we see the archetypical buzz-pollinated flower form (the *Solanum-*type) potentially experiencing a pollinator shift with apparently no phenotypic transition. Further investigation is needed to determine if this indeed represents a functional transition, the extent floral phenotype has been modified, and the drivers (e.g., floral abundance, abiotic conditions, honeybee plasticity, etc.) of these transitions.

Beyond basic research questions, it should also be kept in mind that silverleaf nightshade is a top-noxious weed that is adversely affecting natural, agricultural-, and urban ecosystems across the world. Moreover, its invasive potential remains high, especially in the context of global change: spatial models simulating climatic suitability for this plant indicate numerous currently invadable regions in all continents, with poleward expansions predicted in a warmer atmosphere (Kriticos et al. 2010). Given that honeybees occur with generally high population densities in almost all continents, their impact on the realized fitness of invasive populations of *S. elaeagnifolium*, and the broader implications for natural communities should be explored.

## Acknowledgements

GN and TP would like to thank Chrysoula Tananaki and Georgios Goras for support during fieldwork.

## Conflict of interest statement

The Authors declare no conflict of interest.

## Author contributions

MCM, CMDM, and RRK designed the study. RRK, HL, TP, and GN performed field observations. RRK performed scan microscopy. AK processed the observations, curated media, and wrote the first draft. All co-authors revised the manuscript.

## Electronic Supplementary Material

**Video S1.** A honeybee worker attempting to extract pollen from the flowers of silverleaf nightshade by biting the anthers. Videographer: Georgios Nakas.

**Video S2.** A honeybee worker attempting to extract pollen from the flowers of silverleaf nightshade by holding the anthers with their forelegs and inserting their proboscis through the pores of the anthers (“milking” behavior). Videographer: Georgios Nakas.

**Video S3.** A honeybee worker attempting to extract pollen from the flowers of silverleaf nightshade by shaking the anthers. Videographer: Georgios Nakas.

## Notes

### Competing Interest Statement

The authors have declared no competing interest.

### Summary of Updates

There is a methodology section, a new table, and a map.

